# Experimental evolution at high temperature in a Lepidopteran host of an entomopathogenic nematode and its associated microbiota

**DOI:** 10.64898/2026.09.15.751662

**Authors:** Nell Foucher, Chloé Roux, Jean-Claude Ogier, Stéphanie Bedhomme, Julien Brillard

## Abstract

<u>E</u>ntomo<u>p</u>athogenic <u>N</u>ematodes (EPNs) have a great potential to control various insect crop pests, but their ability to perform their entire life-cycle in insects depends on environmental conditions. The insect killing ability of the endosymbiotic bacterial species (i.e., *Xenorhabdus*) located in the EPNs’ gut (*Steinernema*) has been described for years. More recently, experimental data have suggested that other bacteria species of the nematode microbiota may also contribute to the parasitic success of the EPNs. EPNs and their associated bacteria should therefore be considered as holobionts. Global change, among its many predicted consequences, is expected to increase temperatures which may impair the parasitic success of these EPNs in the insect. Furthermore, higher environmental temperatures could modify the composition of the EPN-associated microbiota and could therefore also contribute to parasitic impairement.

Using a *S. carpocapsae* strain originating from a French orchard, we investigated these hypotheses by experimentally evolving independent EPNs lineages *in vivo* in the lepidopteran species *Galleria mellonella*, at elevated and control temperatures. Several life-history traits of the evolved lineages were measured (insect killing ability, EPN emergence rate, EPN reproduction rate), and the bacterial microbiota of the evolved lineages was characterized by a metabarcoding approach.

The EPNs’ lineages evolved at higher temperature showed impaired emergence rate and their reproduction rate was lower, when compared to lineages evolved at control temperature. The bacterial microbiota composition of these lineages was modified, with particular enrichment in bacteria of the genera *Alcaligenes* and *Pseudomonas*.

Our results provide new evidence that modification of the environment (here, the abiotic parameter temperature) during several successive generations can modify the parasitic success of EPNs, concomitantly to a change in the microbiota composition. A better understanding of the life-history traits of these EPNs should help improve the biocontrol efficiency in a context of global warming.

## Introduction

Entomopathogenic nematodes (EPNs) are naturally found in soils, and several strains of *Heterorhabditis* and *Steinernema* are commercialized as powerful and environmentally friendly agents to control insect pest populations [1–3]. EPNs form a mutualistic symbiosis with bacteria belonging to genus *Xenorhabdus* and *Photorhabdus* of the *Morganellaceae* family. Upon infection of an insect host by the nematode infective juveniles (IJs) [4], the endosymbiotic bacteria are released from the nematode’s gut to the insect hemocoel where they rapidly proliferate, inducing septicemia and killing the host within 48–72 hours. However, the success of these nematobacterial complexes might not be solely dependent on the specific endobacterial symbionts. By analyzing the microbiota of a *Steinernema carpocapsae* strain, it was recently shown that additional bacterial species frequently associated with these nematobacterial complexes, such as a *Pseudomonas protegens* strain with entomopathogenic properties, may enhance the parasitic success, alongside *Xenorhabdus* [5]. This striking discovery reframes EPNs as putative holobionts, where the combined action of a pathobiome drives their ecological and agricultural impact [3].

Yet, the stability and functionality of these pathobiomes might be strongly modified by various external factors (biotic or abiotic), since environmental stressors, particularly temperature fluctuations, can dramatically reshape microbial communities [6]. While EPNs are used for their biocontrol potential, their efficacy in the field is variable, often hinging on temperature [7–10]. For instance, *Heterorhabditis* strains achieve their highest insect mortality rates at 20–25°C. Whereas *Steinernema* thrives at cooler temperatures, it was described as still effective at 30°C but with reduced insect mortality rates [7]. Other studies showed that some EPN strains exhibit enhanced reproductive success when the temperature rises from 23 to 30°C [8]. Temperature-dependent changes in EPN behavior have also been reported, with *Steinernema* being attracted to insect-emitted organic compounds when produced at 15 °C, but repelled when produced at higher temperatures (25 °C) [10]. In addition, some EPN species become less motile and alter their host-seeking behavior under thermal stress [10]. All these phenotypic shifts underscore the remarkable diversity within EPN species, but also raise critical questions: Can these organisms adapt to temperature increase and does their microbiota play a role in this process?

In an era of rapid global change, nematobacterial complexes are likely to face challenging environmental conditions. Assessing a global warming scenario on a microcosm associating a plant, an insect pest and EPNs, showed that elevated temperatures and CO_2_ levels can reduce EPN infestation rates by 40%, putatively exacerbating plant damages by insect pests [9]. Despite the clear link between environmental conditions and EPN performance, no study has explored whether shifts in microbiota composition correlate with these phenotypic changes. Unraveling this connection could improve our knowledge of the potential mechanisms by which EPNs—and their microbial partners— face environmental changes, offering a pathway to bolster their biocontrol potential in a warming world.

Experimental evolution provides a powerful approach to address these questions. Serial passage experiments have revealed the adaptive strategies in various organisms facing abiotic stressors [11] in which the monitoring of molecular and phenotypic evolution provides a better understanding of the causes or consequences of parasite evolution. They have often led to the discovery of new mechanisms potentially involved in the short-term evolution of these organisms [12, 13]. These approaches also provide data that can ultimately be used to test fundamental biological hypotheses, notably concerning how these organisms respond to environmental changes [12, 14]. However, most studies have focused on simplified systems— isolated microbes or dual-species cultures [15, 16], —leaving a gap in our understanding of how complex, multi-species communities [17, 18], evolve in response to environmental pressures. It was shown that response to abiotic stress of a given bacterial species can depend on the presence of other bacterial species in the community [19], revealing the importance of a bacterial community equilibrium. In addition, there are several examples showing the role of microbiomes in protecting their hosts from high temperatures, but these studies often focus on a single species (a mutualistic microorganism) that has co-evolved with its host [20]. While bacterial symbionts can shape host evolution [21, 22], to date few experimental evolution studies have been carried out on more complex systems, coupling host phenotypic changes with shifts in their natural microbial consortia [23], as suggested for *Drosophila*, where the microbiota may be an agent of selection that drives adaptation in the wild [24].

By subjecting a *S. carpocapsae* EPN strain to serial passages *in vivo* at both a control temperature and a challenging high temperature, this study aims to determine if (i) life-history trait of EPNs evolve under thermal stress and (ii) if their bacterial microbiota composition undergo parallel changes. The gained knowledge may improve the strategies of EPNs use in biocontrol approaches in a context of climate change.

## Methods

### Strains and raising conditions

The nematode strain FRA241 of *Steinernema carpocapsae*, was originally isolated in 2017 from a French apple orchard soil in South of France [5], using a classical ex situ *Galleria mellonella* trap [25]. Laboratory-reared EPN stocks were multiplied every nine-ten months by exposing a batch of ten *Galleria* larvae to 1000-5000 IJs (infective juveniles), incubated at 23°C and harvested 21-28 days later following the White trap method [26]. Newly emerged IJs were rinsed on a 20-µm sieve and stored in fresh Ringer’s solution (Merck) at 9°C as previously described [27]. The strain FRA241 used as the nematode ancestral population of this study had therefore been maintained in the laboratory for 5 replicative cycles since its harvest from the field and before its use in the experimental evolution approach.

Insect larvae of *G. mellonella* were reared on an artificial diet (based on honey and wax) and incubated at 28 °C in the dark. Last-instar larvae were collected 28-30 days after oviposition and used for infestation by nematodes.

### Experimental evolution approach by serial passages

12 technical replicates of 10000 infective juveniles (IJ) of the ancestral population of strain FRA241 were prepared. Each of them was used to infest a batch of 10 *Galleria* larvae.

Infestations were performed in Petri dishes layered with Whatmann paper. This allowed the insect larvae to freely move and therefore required that the host-seeking behavior of the EPN was efficient for a proper infestation.

Six of these batches were placed in the dark at the control temperature, hereafter referred as “EE23” (i.e. 23°C, the standard temperature used in the laboratory to reproduce the EPN stock), and 6 were placed at a challenging high temperature hereafter referred as “EE28” (i.e. 28°C). This second temperature was chosen because our preliminary tests showed that the EPN strain used (FRA241) was unable to perform its complete life cycle in *G. mellonella* at higher temperatures (30°C), although it was still able to kill insects’ larvae. Except for the temperature difference, the same standard laboratory conditions were maintained in both treatments.

As a control experiment to verify that insect death was solely due to EPN infestation, batches of 5-10 uninfested *G. mellonella* larvae for each temperature were placed in the same conditions as for the experimental evolution. Three days post infestation, control larvae were always alive, and all dead larvae of the experimental evolution were collected and placed in a White-trap as described above.

At each passage, 14 days post infestation, IJs that emerged from the batch of cadaver were collected, passed through a 20-µm mesh sieve, washed briefly with water and then thoroughly rinsed with sterile Ringer buffer as previously described [27]. They were then quantified by counting serial dilutions under the stereomicroscope, and 10000 freshly harvested IJs were used to infest a new batch of 10 insect larvae, as described above. The experimental design is illustrated in SuppData S1.

The serial passage experiment was launched in August 2022 and was stopped after the 20^th^ serial passage *in vivo*. Based on previous estimations of the number of generations performed in an insect larva [28], our experimental setup represents between 40 and 60 generations of nematodes. During the course of the experimental evolution, in some lineages the infestation sometimes did not lead to emergence, or it led to a drastically low number of IJs. In such cases, the infestation cycle had to be repeated. Consequently, the whole experimental evolution lasted for 10 to 18 months, depending on the lineages. When each of the evolved lineages reached their 20^th^ serial passages *in vivo*, the emerging nematodes (IJs) were harvested, sieved (20 µM), washed and then stored in Ringer buffer at low temperature (9°C) for long-time conservation. In these conditions, they could stay alive 9-12 months until a new insect infestation would be performed.

Because the 12 evolved lineages reached their 20^th^ serial passages *in vivo* at different times, the time they spent in storage condition at low temperature (i.e., in Ringer at 9°C) varied and this parameter was suspected to have an impact on their fitness. To avoid this potential confounding factor, we performed an additional cycle *in vivo* just before the life-history trait and physiological characterization of evolved lineages. We used the exact same conditions as during the serial passage experiment, to allow all the evolved lineages to be synchronized and freshly harvested.

### Life-history traits assays

Three parameters of the life-history traits of evolved nematode lineages were monitored, (i) ability to kill the insect host, (ii) ability to perform a complete life cycle in this host, (iii) total number of IJs emerging from the cadaver, as previously described [27] and detailed below. These parameters can be used as proxis to estimate the overall fitness of each of the evolved lineages.

In order to systematically evaluate the effect of the temperature on each of these three parameters, insect infestations were carried out at both the control (i.e. “C” = 23°C), and the challenge temperatures (i.e. “C+5” = 28°C), for all lineages whatever their experimental evolution temperatures.

Briefly, for each lineage, twenty *G. melonella* larvae were individually infested by 100 IJs and then incubated in the dark, at 23°C for 10 of them, and at 28°C for the other 10. Three independent replicates were performed (totalizing 30 insect larvae for each nematode lineage and for each temperature tested). In parallel, twelve independent replicates of the ancestral lineage were also tested in the same conditions (6 replicates at 23°C and 6 at 28°C) as a control.

Then, (i) Insect survival was monitored for each nematode lineage after 48h, and up to 72h after infestation.

(ii) Dead larvae were placed individually on White traps [26]. After 14, 21 and 28 days, the parasitic success (defined as emergence of nematodes from the insect cadavers) was monitored.

(iii) The reproductive success (defined as the number of IJs that emerged from each insect larvae) was quantified under the stereomicroscope as described above, at 14, 21 and 28 days post-infestation.

Mortality data were analysed using generalized linear mixed models with logit link function. Emergence rate data were analysed with a generalized linear mixed model with logit link function. The reproductive success data were analysed with a generalized linear mixed model fitted with a negative binomial distribution to account for overdispersion. The details of each model and how they were compared and chosen is given in the result section.

### Characterization of the Microbiota associated to nematodes lineages

EPN’s bacterial microbiota composition was analyzed by a metabarcoding approach using protocols developed in our laboratory, as previously described [29] and detailed below.

### DNA extraction

Prior to DNA extraction, infective juveniles (IJs) of nematodes were collected after one cycle and from evolved lineages when they reached their 20^th^ cycle of EE. For each lineage, 2 to 3 replicate samples were used. Each sample was constituted of 1,000-5,000 IJs and was placed into a 2 ml screw-cap tube compatible with bead-beating. Samples were subjected to three washing cycles (2,000 rpm for 1 min) with Ringer’s solution to remove bacteria present in the surrounding medium and on the nematode surface. DNA extraction was then performed as previously described [5]. Briefly, samples were subjected to mechanical and enzymatic lysis, followed by proteinase K treatment, RNase digestion, and DNA purification steps as previously described. Negative extraction controls were included to monitor potential contamination.

### Library preparation and Illumina MiSeq sequencing

Amplicon libraries targeting the bacterial *rpoB* gene were prepared using a nested PCR approach as previously described [30]. Briefly, the first PCR (25 cycles) amplified an approximately 900 bp fragment of the *rpoB* gene, followed by a second PCR (15 cycles) generating an approximately 520 bp amplicon containing Illumina adapter overhangs. Negative and positive controls were included at each amplification step to monitor contamination and amplification efficiency. Amplicons of the expected size were then sent to the Genseq platform (University of Montpellier, France) for purification, indexing, pooling, and sequencing on an Illumina MiSeq platform (2 × 300 bp).

### Bioinformatic processing of sequence data

Sequencing reads were processed using FROGS v4.1.0 [31]. Briefly, the pipeline included successive steps of pre-processing, clustering using Swarm, chimera removal, cluster filtering, and taxonomic assignment. Taxonomic affiliation was performed using the *rpoB* reference database available at https://genoweb.toulouse.inrae.fr/frogs_databanks/assignation/rpoB/. All analyses were conducted using the parameters as previously described [30].

### Bacterial community and statistical analyses

Microbial community analyses were performed using the R package phyloseq [32], vegan [33], Microbiota Process and DESeq2 [34]. We used ggplot2 to generate the plots.

#### ASVs Filtering

Amplicon Sequence Variants (ASVs) were filtered by retaining only those representing at least 0.1% of reads per sample to remove artifactual sequences and low-abundance taxa, as recommended [29].

#### Species aggregation

Taxonomic assignments were curated based on RDP bootstrap confidence values: species-level annotations with confidence scores below 0.8 were conservatively reclassified at the genus level, indicated as “sp.”. Conversely, ASVs with identical species-level assignments and bootstrap scores above 0.8 were merged, and their read abundances summed to avoid redundancy.

#### Rarefaction curves

Rarefaction curves were generated using the ggrare function from the ranacapa package in R. Prior to visualization, samples were rarefied to an even sequencing depth of 10,000 reads using the rarefy_even_depth function (phyloseq package), in order to standardize sampling effort across samples. Rarefaction curves were computed with a step size of 100 sequences and without confidence intervals.

#### Taxonomic Composition and Visualization

Sequencing data were transformed into relative abundances by normalizing each sample to its total read count. Barplots were generated at species levels using the get_taxadf function of MicrobiotaProcess.

#### Alpha diversity

We used a two-way analysis of variance (ANOVA) to assess the effects of incubation, cycle, and their interaction on alpha diversity. Although normality was not strictly assumed, ANOVA is robust to moderate deviations from normality for continuous diversity indices. Pairwise Wilcoxon tests were used as post-hoc analyses to explore differences between specific groups.

#### PcoA and Permanova

Beta diversity was assessed using Bray–Curtis dissimilarities calculated on Hellinger-transformed data. Principal Coordinates Analysis (PCoA) was used to visualize patterns of community composition across samples.

Differences in microbial community structure were tested using permutational multivariate analysis of variance (PERMANOVA) based on 9,999 permutations. The model included incubation temperature, the cycle, and their interaction (incubation × cycle).

### Comparisons of relative abundance of taxa at 23°C and 28°C

#### Differential Abundance Analysis (DESeq2)

Prior to differential analysis, low-abundance taxa were filtered to reduce sparsity and improve statistical power: only ASV representing at least 0.1% of reads per sample were retained, as described above. Differential abundance analyses were performed to identify microbial taxa differing between incubation temperatures (23°C and 28°C) using the DESeq2 framework [34]. Amplicon sequencing data were processed as raw count tables within the phyloseq environment in R. Samples were grouped according to incubation temperature, and differential abundance was assessed using a design formula including incubation as the explanatory variable. The reference level was set to 23°C, allowing estimation of log2 fold changes for taxa enriched at 28°C relative to 23°C. Size factors were estimated using a geometric mean approach robust to sparsity, as recommended for microbiome count data containing many zeros. Dispersion parameters were estimated using a parametric fit, and differential abundance testing was conducted using the negative binomial generalized linear model implemented in DESeq2.

Resulting p-values were adjusted for multiple testing using the Benjamini–Hochberg false discovery rate procedure. ASVs with adjusted p-values < 0.05 were considered statistically significant. Taxa with positive log2 fold changes were interpreted as enriched at 28°C, whereas those with negative values were enriched at 23°C.

For visualization, significant taxa were displayed using heatmaps based on normalized counts.

### Relative abundance of selected taxa

Relative abundances of selected taxa (Alcaligenes faecalis, Pseudomonas sp. and Xenorhabdus nematophila) were extracted from the dataset. Differences in relative abundance between incubation temperatures (23°C and 28°C) were assessed using non-parametric Wilcoxon rank-sum tests, as data did not meet assumptions of normality. P-values were adjusted for multiple testing using the Benjamini–Hochberg method when applicable. Graphical representations were generated using ggplot2, displaying boxplots of relative abundances across conditions.

## Results

### Effect of incubation temperature on life-history traits of the ancestral strain

The incubation temperature (23°C vs 28°C) had no effect on the insect mortality rate (measured 2 to 3 dpi) as 59 out of 60 insect larvae died in the 23°C treatment and all 60 insect larvae died in the 28°C treatment. To assess the effect of incubation temperature on the emergence rate (i.e., the number of insect cadavers for which IJs emergence was observed), we used a generalized linear mixed model (glmer function in R), with logit link function and with incubation temperature as fixed effect and block as random effect. The incubation temperature had no effect on emergence at 14 dpi (estimate=-0.087±0.189, *z*=-0.459, *p*=0.646), at 21 dpi and at 28 dpi (estimate=9.249*10^-15^ ± 2.853*10^-1^, *z*=0, *p*=1, for both). Reproductive success data were analysed using a generalized linear mixed model fitted with a negative binomial distribution (nbinom2 family, log link) to account for overdispersion (R package glmmTMB) with incubation temperature as fixed effect and block as a random effect.Temperature had no significant effect on reproductive success at 14 dpi (estimate=0.019±0.038, *z*=0.499, *p*=0.618) but a significant effect on reproductive success at 21 dpi (estimate=-0.123±0.039, *z*=-3.185, *p*=0.001) and at 28 dpi (estimate=-0.281±0.032, *z*=-8.656, *p* <0.001), with more IJs emerging at 23°C than at 28°C at the two time points (**Fig 1**).

**Fig. 1:**
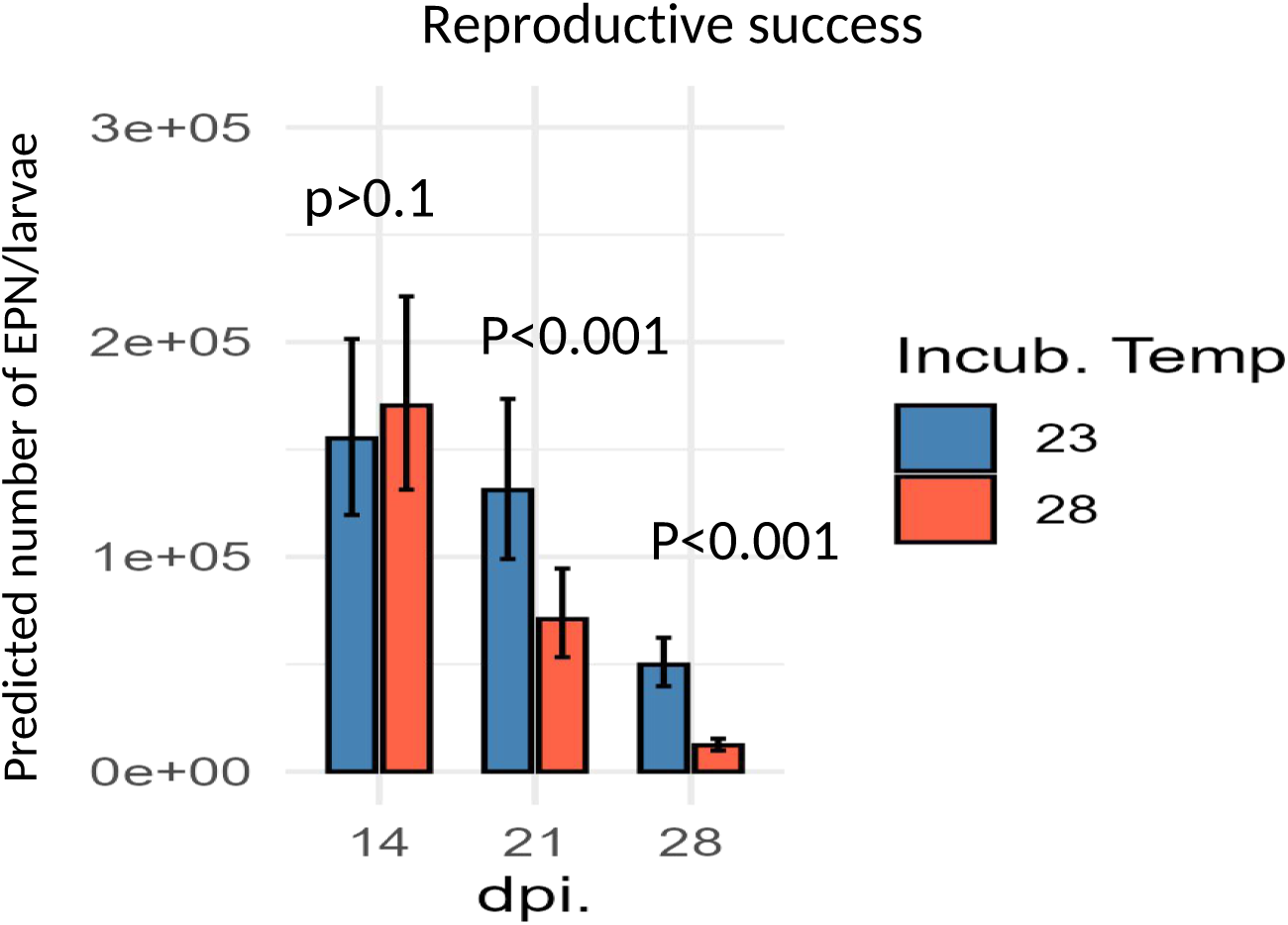
Effect of incubation temperature on the reproductive success of the ancestral strain *Galleria mellonella* larvae were infested by 100 *Steinernema carpocapsae* IJs, from 6 independent replicates of the ancestral FRA241 strain (ancestral), and incubated at either 23°C (blue) or 28°C (red). Dead larvae were put on a White-trap. Emergence of IJs from *G. mellonella* cadaver, as well as the number of total IJs were monitored 14, 21 and 28 days post-infestation (dpi). A Generalized linear mixed model (GLMM), with binomial distribution was applied. Graphs represent the GLMM predicted data of the emergence observed from 10 larvae for each of the 6 replicates of the ancestral strain, per temperature.

### Evolved EPN life-history traits: experimental evolution at high temperature did not change EPN insect killing efficiency

All evolved lineages were assayed at two incubation temperatures (23°C and 28°C). Mortality data were analysed with a generalized linear mixed model (glmer function in R), with logit link function and with evolution temperature and incubation temperature as fixed effect and lineage as random effect. Neither evolution temperature nor incubation temperature had a significant effect on mortality (*z*=-1.105, *p*=0.269 and z=-0.455, p=0.649, respectively), with mortality rates above 0.96 for the four treatment combinations, *i.e.* similar to the ancestral lineage.

These results show that the insect mortality rate caused by EPNs did not change during experimental evolution at low and high temperature and remained unaffected by the incubation temperature.

### Evolution at high temperature induced a decrease in EPN emergence rate and reproductive success

The emergence rate was quantified at 14 dpi, to fit the experimental evolution process, as well as 21 or 28 dpi, to investigate any change in emergence dynamics that may occur in the evolved lineages.

The emergence rate data were analysed with a generalized linear mixed model (glmer function in R), with logit link function and the reproductive success data were analysed with a generalized linear mixed model fitted with a negative binomial distribution (nbinom2 family, log link) to account for overdispersion (R package glmmTMB). For each variable, two models were built: (1) model with evolution temperature, incubation temperature and their interaction as fixed effects and lineage nested within evolution temperature as a random intercept and (2) a reduced version of the previous one without the interaction between evolution temperature and incubation temperature. For each variable and each time point (14, 21 and 28 dpi), the two models were compared by Akaike Information Criterion. The detailed results of this statistical analysis are contained in **table 1**.

**Table 1.**
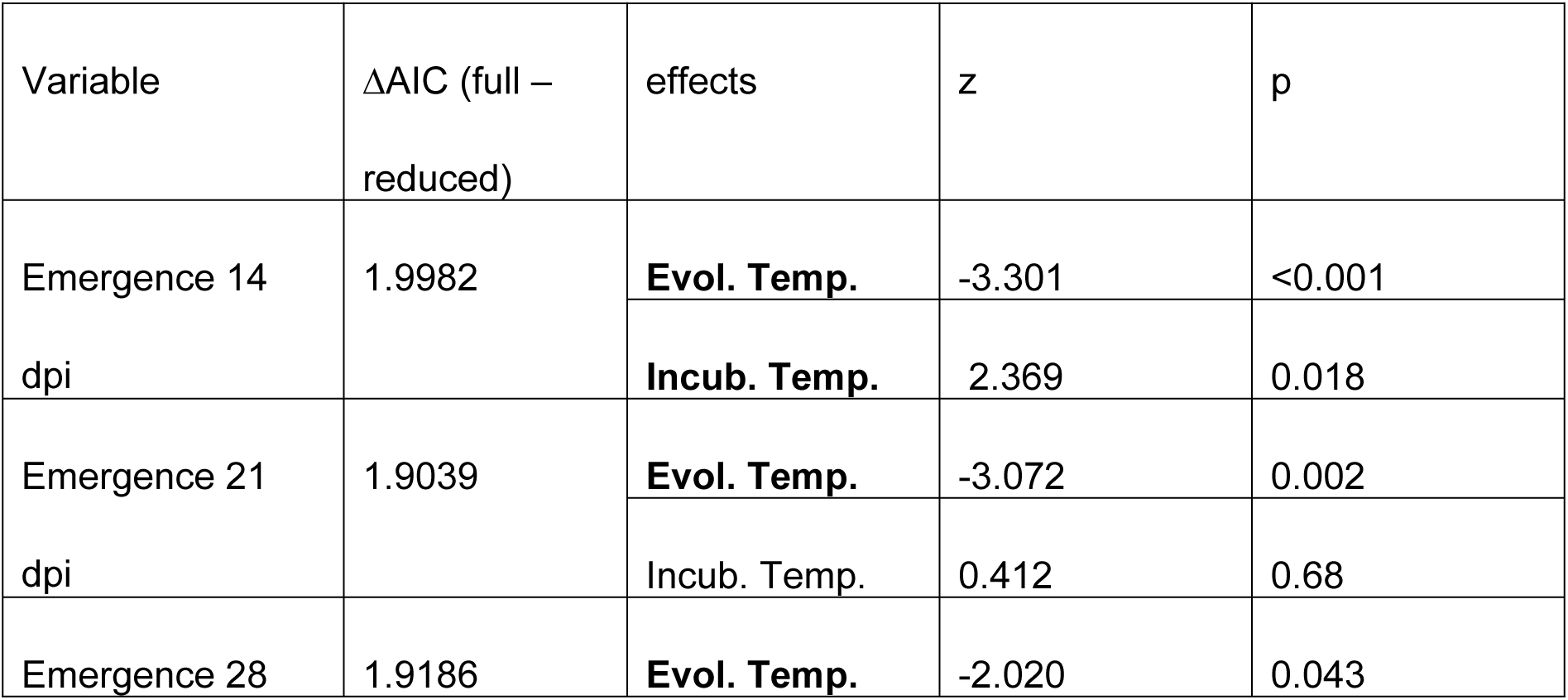
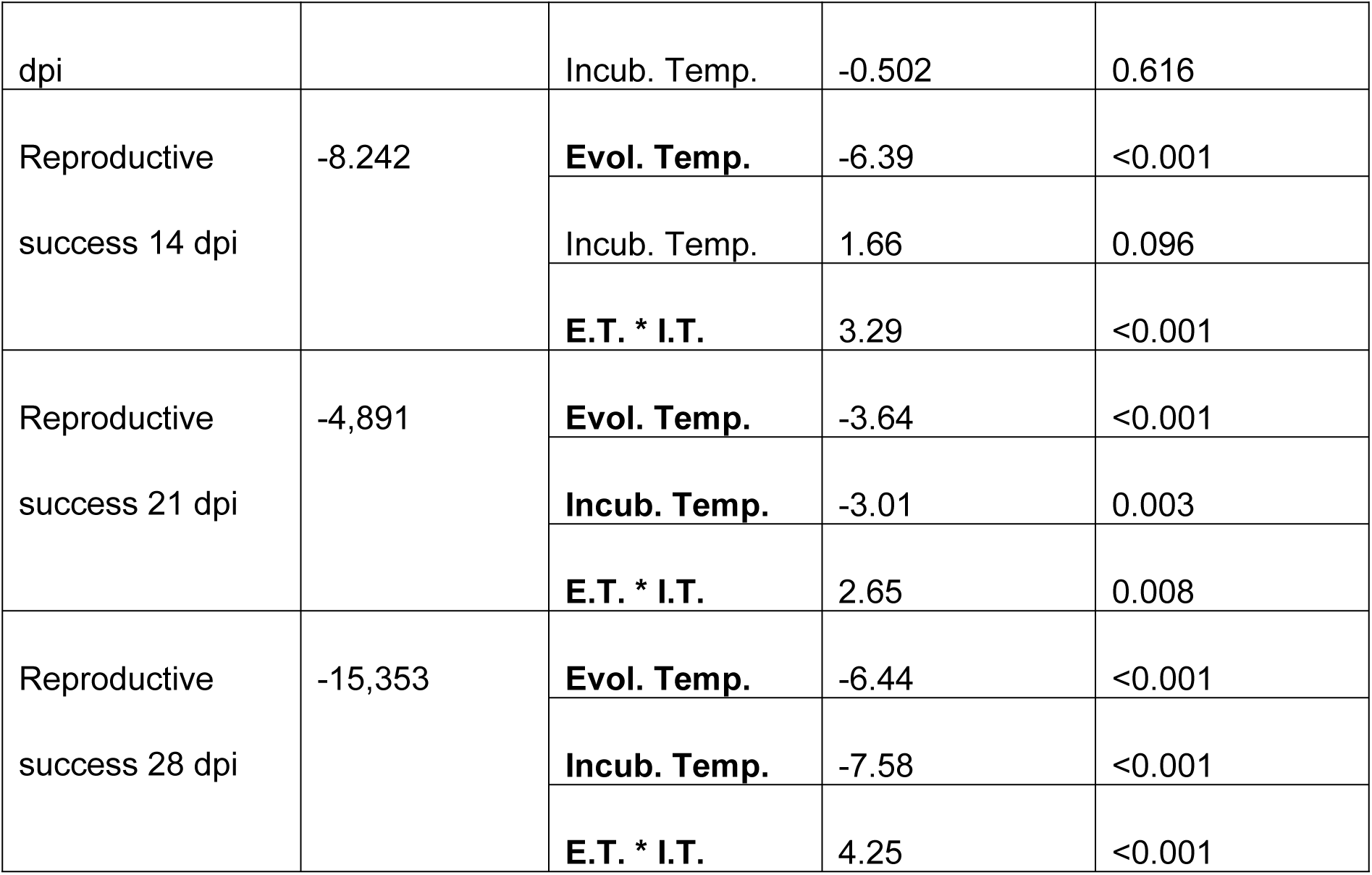

Globally, the temperature of evolution had a significant effect on both variables at all time points, with populations evolved at 28°C having a lower emergence probability (**figure 2**) and a lower reproductive success (**figure 3**) than populations evolved at 23°C. Additionally, for emergence rate at 14 dpi, there was a significant temperature effect with a higher emergence rate for incubation at 28°C (**figure 2**). However, incubation temperature had no significant effect on the emergence rate at 21 and 28 dpi and the interaction between evolution temperature and incubation temperature was never significant. Regarding reproductive success, the interaction between evolution temperature and incubation temperature was significant at the three time points but with different patterns depending on time point. At 14 dpi, there was a higher reproductive success after incubation at 28°C and this was more marked for lineages evolved at 28°C than for lineages evolved at 23°C. At 21 and 28 dpi, there was a higher reproductive success after incubation at 23°C for lineages evolved at 23°C whereas in lineages evolved at 28°C, the reproductive success was equivalent after incubation at 23°C and 28°C. In all cases, emergence rates and even more so reproductive success of evolved lineages were lower than those of the ancestral lineage (**figure 2** and **3**).

**Fig. 2:**
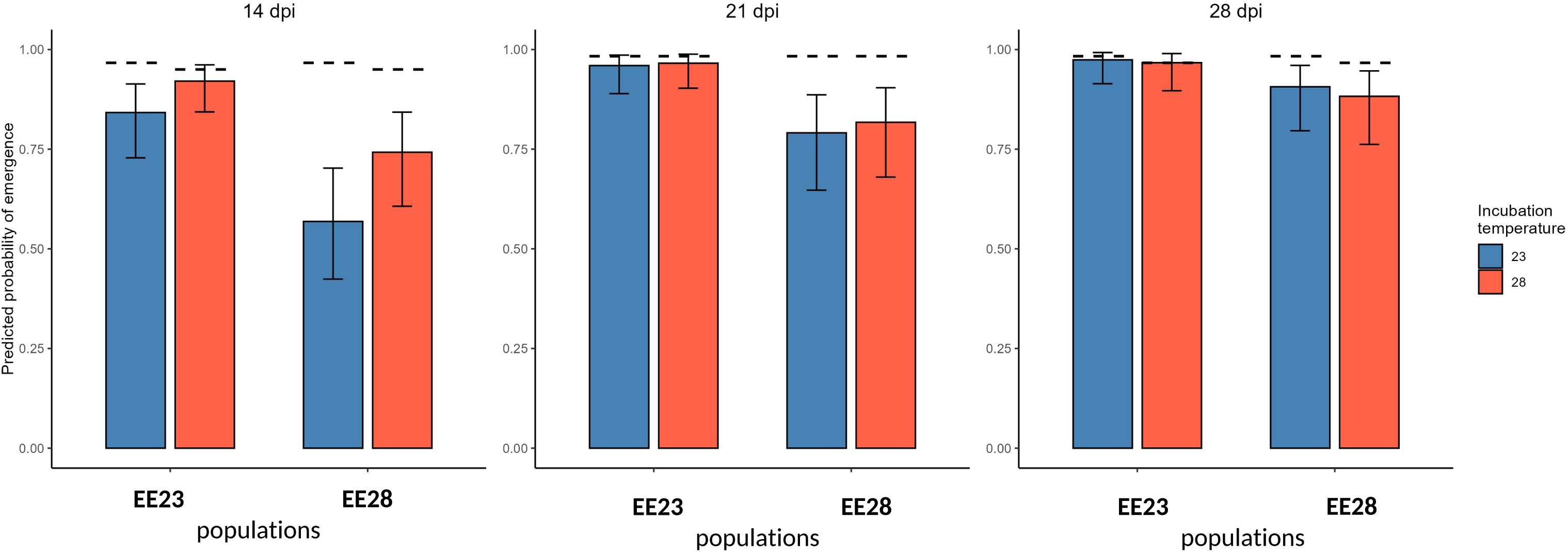
Emergence rate *G. mellonella* larvae were infested by 100 *S. carpocapsae* IJs, from 6 evolved lineages after 20 serial passages at 23°C (EE23) or at 28 °C (EE28), and incubated at either 23°C (blue) or 28°C (red). Dead larvae were put on a White-trap, and emergence of IJs from *G. mellonella* cadaver was monitored 14, 21 and 28 days post-infestation (dpi). A Generalized linear mixed model (GLMM), with binomial distribution was applied. Graphs represent the GLMM predicted probability of emergence of the 12 evolved lineages after serial passages at the control temperature (6 lineages, EE23), and at the challenge temperature (6 lineages, EE28), each of them tested at both the control (23°C) and the challenging (28°C) temperatures. Each experiment contained 10 larvae per lineage/temperature (totalizing 60 larvae for each of the presented histogram). Dashed lines represent the emergence rate observed for the ancestral lineage tested in the same conditions.

**Fig. 3:**
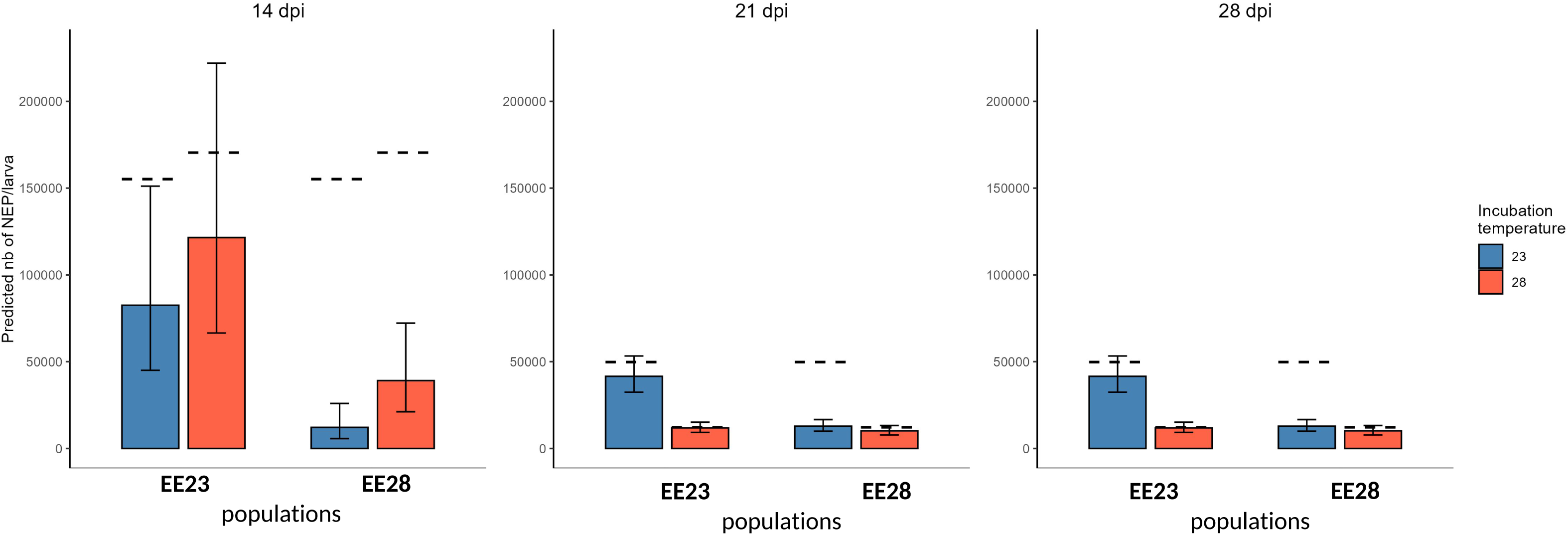
Reproductive success *G. mellonella* larvae were infested by 100 *S. carpocapsae* IJs, from 6 evolved lineages after 20 serial passages at 23°C (EE23) or 6 evolved lineages at 28 °C (EE28), and incubated at either 23°C (blue) or 28°C (red). The dead larvae were put on a White-trap and incubated at either 23°C (blue) or 28°C (red). IJs emerging from *G. mellonella* cadavers were harvested 14, 21 and 28 days post-infestation (dpi) and the number of total IJs was monitored. A Generalized linear mixed model (GLMM), with binomial distribution was applied. Graphs represent the GLMM predicted data of the number of IJs emerging from each cadaver. Each experiment contained 10 larvae per lineage/temperature (totalizing 60 larvae for each of the presented histogram). Dashed lines represent the number of IJs emerging for the ancestral lineage tested in the same conditions.

Because the incubation temperature had a significant effect on the reproductive success but with different patterns depending on time point, we analyzed the cumulated data of IJs that have emerged at the end of the experiment, i.e., the reproductive success at 14+21+28 dpi (NB: with this EPN strain no emergence is observed later than 28 dpi). Results (SuppData S2) indicate that altogether, the total amount of IJs from EE28 were lower than those from EE23, whatever the tested temperature.

### Effect of the temperature on EPN bacterial microbiota composition before and after experimental evolution

The composition of the bacterial microbiota associated to the EPNs evolved lineages was analyzed by a metabarcoding approach using the *rpoB* marker, a method previously shown to be highly efficient for EPNs [5]. We also performed metabarcoding analyses on a reference lineage prior to the experimental evolution. To avoid potential bias due to prolonged storage of the original strain at 9 °C, the first passage of strain FRA241 in *Galleria* larvae (cycle 1, incubated at either 23°C or 28°C) was used as the ancestral reference lineage.

Before the filtration step, a total of 2,139,642 raw sequences were obtained from 65 samples, corresponding to 37 and 28 replicates for cycles 1 and 20, respectively, representing 386 unique amplicon sequences across all individual samples (see SuppData S3 - Table ASV).

After applying a 0.1% filtration threshold and aggregating amplicon sequences corresponding to the same species (see Materials & Methods for details), the number of ASVs (Amplicon Sequence Variants) decreased to 148. Richness rarefaction curves (Supplemental Data S4), generated separately for evolution cycles 1, and 20, reached a saturation plateau, indicating that the sampling effort and sequencing depth were sufficient to capture the majority of microbial species.

The taxonomic compositions of the microbiota associated with the ancestral reference lineage (cycle 1) and the experimental evolution (cycle 20) at 23 °C and 28 °C are presented in Supplementary data S5. Overall, the EPN-associated microbiota was largely dominated by the phylum Pseudomonadota, while only a minor fraction of ASVs was affiliated with other phyla, including Bacteroidota and Bacillota (**Supp Data S5.A).**

Subsequent analyses focused on the genus (**Supp Data S5.**B) and species-level (**Supp Data S5.**C), with the latter enabled by the *rpoB* marker, which provides reliable species-level taxonomic assignment.

In the reference lineage, four genera predominated: *Achromobacter, Xenorhabdus* (the obligate symbiotic bacterial partner of EPNs), *Brevundimonas*, and *Serratia*. These taxa have previously been reported as members of the so-called “second bacterial circle” associated with EPNs [3].

Microbiota associated with the evolved lineages were largely dominated by *Alcaligenes*, particularly under the high-temperature challenge, followed to a lesser extent by *Xenorhabdus*.

Several additional genera—namely *Pseudomonas*, *Brevundimonas*, *Pseudochrobactrum*, *Delftia*, *Stenotrophomonas, Devosia, Achromobacter, Comamonas*, and *Brucella*—were consistently detected across samples but occurred at lower relative abundances. At the species level (**Supp Data S5.**C), the most abundant taxa included *Xenorhabdus nematophila*, *Pseudomonas solani, Brevundimonas nasdae, B. diminuta, Alcaligenes faecalis, Stenotrophomonas maltophilia*, and *Delftia acidovorans*.

We then estimated the alpha-diversity of the bacterial microbiota using the Observed and Shannon indexes (**Fig**. **4A**). The results indicate that the experimental evolution temperature had a significant effect on alpha-diversity, as evidenced by significant changes in both the Observed and Shannon indices. We observed a lower bacterial richness and diversity for lineages evolved at 28°C compared to lineages evolved at 23°C.

**Fig. 4:**
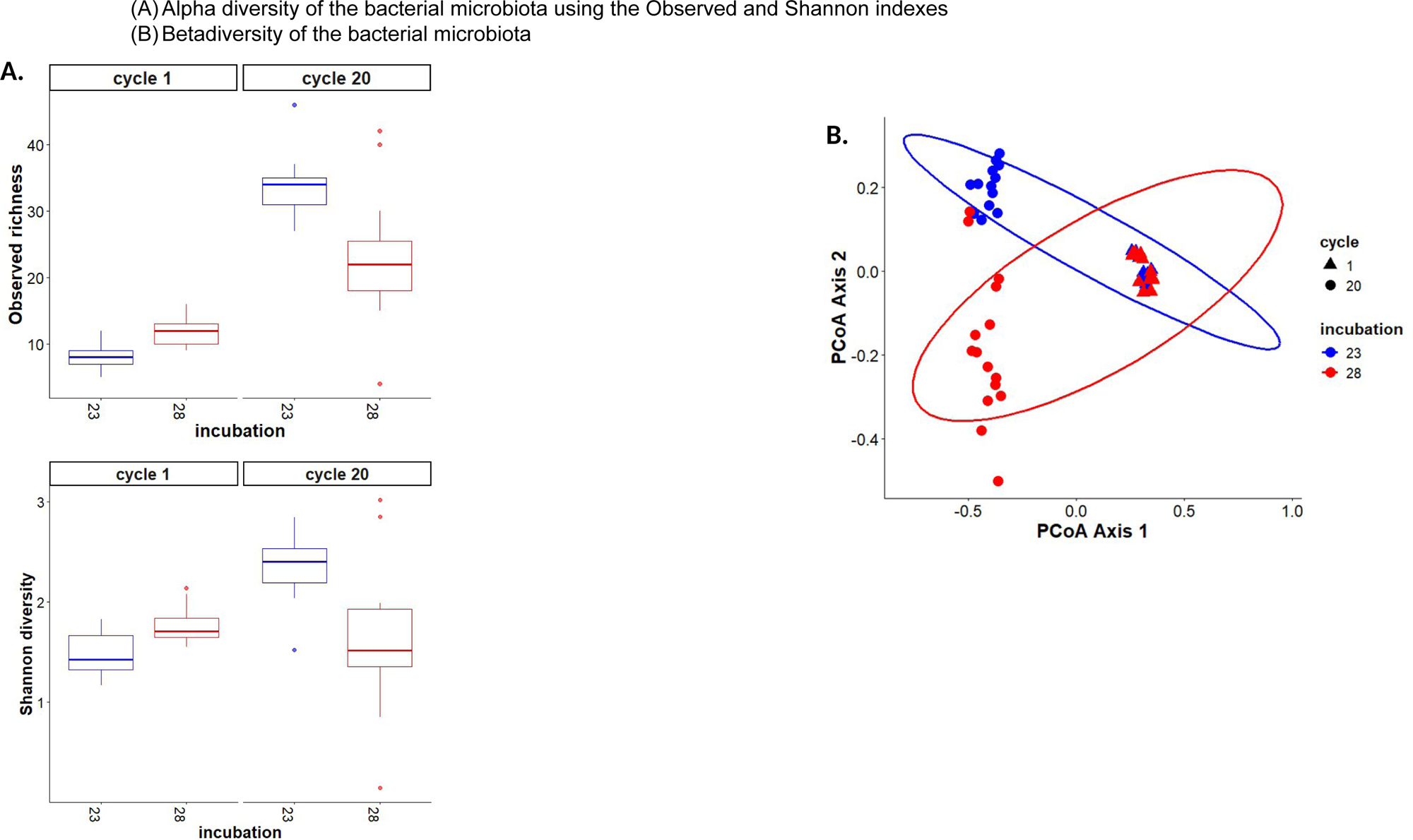
Characterization of the EPN microbiota. **(A) Alpha diversity of the bacterial microbiota using the Observed and Shannon indexes** Alpha diversity responded differently depending on the metric considered. Observed richness was significantly affected by both incubation temperature and evolutionary cycle, with a strong effect of cycle (F = 179.68, p < 2 × 10^−16^) and a significant incubation × cycle interaction (F = 31.32, p = 5.51 × 10^−7^), indicating that temperature effects depended on evolutionary history. In contrast, Shannon diversity showed no significant main effect of incubation, but was significantly influenced by cycle (F = 11.56, p = 0.00119) and displayed a strong interaction between incubation and cycle (F = 25.67, p = 4.02 × 10^−6^). Post-hoc Wilcoxon tests (BH-adjusted) confirmed that most pairwise comparisons between incubation × cycle groups were significant for both richness and Shannon diversity (p < 0.01), supporting the presence of strong non-additive effects of incubation and evolutionary cycle on microbial alpha diversity. **(B) Betadiversity of the bacterial microbiota** PERMANOVA based on Bray–Curtis distances revealed that bacterial community composition was strongly structured by experimental factors. The model including incubation and cycle explained 64% of the total variance (R^2^ = 0.64, F = 55.41, p = 0.0001). When the interaction term (incubation × cycle) was included, the model explained 69% of the variance (R^2^ = 0.69, F = 46.10, p = 0.0001), indicating that the effect of incubation on community composition depends on the evolutionary cycle.

We further investigated beta-diversity among the two temperature treatments at cycles 1 and 20 using Principal Coordinates Analysis (PCoA) based on Bray–Curtis distances (**Fig. 4B**). The PCoA plots revealed two distinct clusters corresponding to lineages evolved at the control (23 °C) and challenged (28 °C) temperatures. Significant differences in microbiota composition between the two temperature regimes were confirmed by PERMANOVA (**Fig. 4B**) for cycle 20. The proportion of variation explained by temperature (R^2^) was 28% for cycle 20, indicating that temperature influences microbiota composition at the end of experimental evolution. In contrast, data at cycle 1 clustered regardless of the temperature, indicating that temperature had no influence on microbiota composition after just one cycle.

To obtain a more detailed assessment of changes in bacterial composition among the evolved lineages, differentially abundant ASVs between the two evolution temperatures were identified using DESeq2. DESeq2 results are visualized as a heatmap (**Fig. 5**), with log2 fold-change values reported in Supp Data S3. The heatmap displays ASVs showing significant differential abundance (adjusted p-value < 0.05) between lineages evolved at 23 °C (EE23) and 28 °C (EE28).

**Fig. 5:**
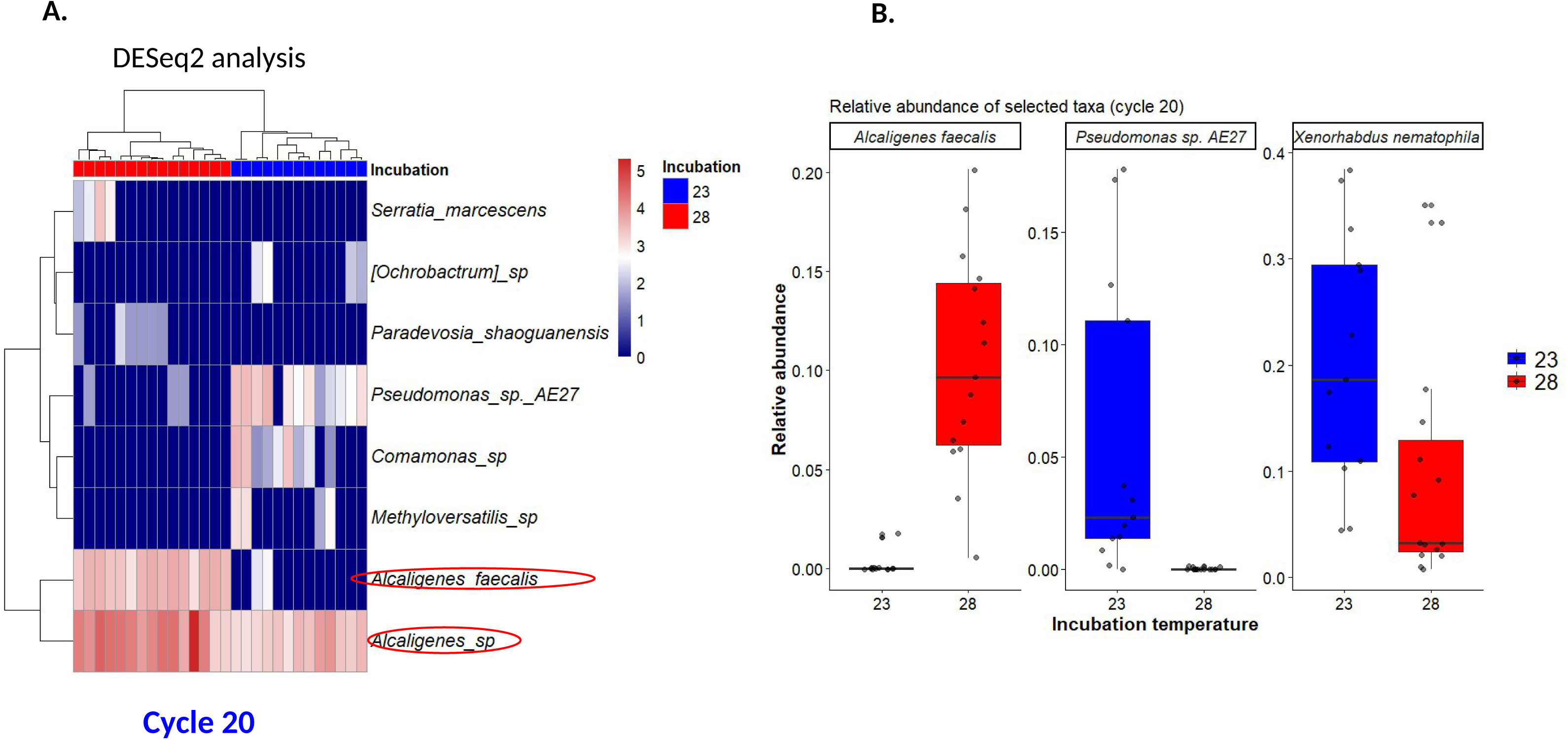
Relative abundance of some of the EPN microbiota bacterial taxons. **(A) Differentially abundant ASVs between the two incubation temperatures identified using DESeq2a** **(B) relative abundances of *A. faecalis, Pseudomonas* sp. AE27 and *X. nematophila* after the evolution experiment** The relative abundances were compared between incubation temperatures using Wilcoxon tests, and revealed significant differences between evolution temperatures for the three selected taxa

Among the differentially abundant ASVs, the most pronounced changes at the high-temperature regime (28 °C) involved taxa belonging to Alcaligenes. Two taxa, *Alcaligenes* sp. and *Alcaligenes faecalis*, were enriched in lineages evolved at 28 °C compared with the control temperature (23 °C). Conversely, *Pseudomonas* sp. AE27 displayed higher abundance in EE23 lineages, indicating distinct temperature-associated abundance patterns (Fig. 5).

To further characterize the abundance patterns of taxa highlighted by DESeq2, relative abundances of *A. faecalis* and *Pseudomonas* sp. AE27 were compared between incubation temperatures within cycle 20 using Wilcoxon tests. In addition, the symbiotic bacteria *X. nematophila* was included as a reference taxon to assess its specific response to temperature variation. Relative abundance comparisons within cycle 20 revealed significant differences between incubation temperatures for the three selected taxa. *Alcaligenes faecalis* displayed a marked increase at 28 °C (adjusted p = 1.91 × 10^−5^), whereas *Pseudomonas* sp. AE27 and *X. nematophila* showed lower relative abundances at 28 °C (adjusted p = 2.91 × 10^−5^ and 8.29 × 10^−3^, respectively). These results confirm taxon-specific responses to temperature, with contrasting abundance shifts among bacterial taxa during experimental evolution.

Together, these results indicate that experimental evolution temperature shaped the relative abundance of specific bacterial taxa, with *Alcaligenes* taxa being associated with the 28 °C condition and *Pseudomonas* sp. and *X. nematophila* showing reduced abundance under this temperature regime.

## Discussion

### Role of Temperature in EPN Evolution

Experimental evolution provides a powerful tool to investigate how organisms adapt to environmental pressures [11]. In the present study, we performed experimental evolution of a *S. carpocapsae* strain at control and elevated temperatures using *Galleria mellonellla* as insect host and characterised the EPN life-history traits and the composition of the nematode microbiota of the experimentally evolved lineages. The goal was to determine whether and how the EPN could adapt to an elevated temperature and the potential role of its associated microbiota in this phenomenon.

The role of the EPN symbiotic bacteria, *Xenorhabdus* for *Steinernema*, in insect mortality is well-documented in our biological model as well as in the *Photorhabdus-Heterorabditis* system [35, 36]. The timescale of our experiment aligns with established literature on phenotypic adaptation. Previous studies have shown that nematodes can exhibit modified traits within as few as 30 generations [28, 37], while their bacterial symbionts, such as *Xenorhabdus*, may require up to 200 generations to display significant changes [38]. Over the course of the 20 successive cycles performed here, the nematodes have undergone 40 to 60 generations, and *Xenorhabdus* should have been through up to 300 generations exclusively *in vivo*.

Life-history traits measured on the ancestral strain at 23° and 28°C did not reveal any difference induced by the temperature treatment on the mortality, on the emergence rate and on the reproductive success at 14 dpi. The elevated temperature resulted in lower reproductive success at 21 and 28 dpi. This means that the elevated temperature was indeed stressful as it reduced the total reproductive success but in the experimental evolution protocol, the IJs used to start a new cycle of infection were harvested at 14dpi, i.e. at a time point of the life cycle at which the stress imposed by the elevated temperature is not expressed. This choice of harvesting time allowed to perform more cycles of experimental evolution but placed us in sub-optimal conditions to observe adaptation to the elevated temperature conditions. The characterization of the life-history traits of the experimentally evolved lineages actually did not reveal any pattern of adaptation to the elevated temperature which would have been a better performance at 28°C of the lineages evolved at 28°C than the lineages evolved at 23°C. For some traits, such as the emergence rate and the reproductive success at 14 dpi, lineages evolved at 28°C performed better at 28°C than at 23°C but they never performed better than the lineages evolved at 23°C and measured at 28°C. A striking result was that experimental evolution lineages, evolved either at 23°C or at 28°C had reduced emergence rate and reproductive success at all time point after infection than the ancestral line. This could be due to genetic drift, typically caused by a small number of individuals founding each generation and potentially to low initial genetic diversity in the *Steinernema* ancestral population. There are no data on the genetic diversity of the ancestral population and it is difficult to evaluate the effective population size in the experimentally evolving lineages. Indeed, the actual number of founders per generation is uncertain: while each group of 10 insect larva was exposed to 10000 IJs, we do not know how many entered each larva and how many contributed to the sexual reproduction for the production of the next generation of EPNs. However, the reproductive success is significantly lower for lineages evolved at high temperature, suggesting that drift has stronger effects at high temperature, a phenomenon that could participate in EPN fitness decline with temperature elevation.

### Temperature as a Driver of Microbial Diversity

Phenotypic data on EPN microbiota remain scarce, but it is reasonable to hypothesize that different bacterial species within the microbiota possess distinct temperature optima. Temperature can affect the fitness of different microbiome species in distinct ways, both through their individual temperature optima and via altered species interactions. These ecological effects can drive rapid shifts in microbiome composition—even within a single cycle—and may either buffer the host from abiotic stress or exacerbate it, for example if a key microbial function is lost due to the decline of a temperature-sensitive species.

Our finding revealed that the composition of the microbiota is not static but is altered after 20 successive infection cycles in a different way depending on the temperature regime. We observed a lower richness and diversity under elevated temperature, as well as pronounced shifts in overall community structure when EPNs are maintained at 28 °C compared to 23 °C. The observed changes in abundance of some bacterial species composing the microbiota may affect the EPN life history traits and could consequently drive its evolution as described for other organisms [39, 40].

Temperature is a pivotal abiotic factor reshaping microbial communities, as evidenced across diverse host systems [6]. For instance, in the gut microbiota of a beetle (*Zygogramma bicolorata)*, temperature treatments significantly altered alpha diversity in only one generation, demonstrating how thermal regimes can restructure microbial assemblages [41]. In wild flies (*Drosophila melanogaster)*, the daily maximum temperature covaried with the microbiota, suggesting that this factor is a major driver of variation in microbiota composition [42]. Similarly, studies on *Yersinia* in fleas revealed temperature-dependent shifts in bacterial virulence and transmission dynamics, illustrating the intricate link between temperature, microbiota, and host fitness [43]. In addition, it was shown that the microbiomes can play a role in protecting some hosts from abiotic stresses, such as high temperatures [44].

### The Role of Microbiota in EPN Life-History Traits

The presence of specific bacteria can influence nematode life-history traits, such as growth rates, fecundity, and lifespan. Studies have shown that *Caenorhabditis elegans* raised in axenic conditions exhibit developmental delays and reduced reproductive success, highlighting the microbiota’s role in normal physiology [40, 45]. Despite the described critical role of the endosymbiotic bacteria (*Xenorhabdus* for *Steinernema* or *Photorhabdus* for *Heterorabditis*) in EPNs’ fitness [35], the importance of the other frequently associated bacterial species members of the microbiota remains understudied [3] but our findings show a concomitant change in bacterial communities and EPNs’ fitness .

The hologenome theory [46] posits that the host and its microbiota function as a single evolutionary unit. However, not all interactions are beneficial; some microbiota may exploit their hosts, leading to conflicts, and trade-offs between lifespan and fecundity may occur [47]. Distinguishing true co-evolution from transient associations remains a challenge, but evidence from *Drosophila* suggests that even modest shifts in microbiomes can alter host populations [24]. Our study provides preliminary evidence that serial passaging at sub-optimal temperatures alters EPN life-history traits and modify microbiota composition, suggesting a potential feedback loop between host evolution and microbial dynamics. Whether the microbiome affects directly the EPN evolutionary potential by shifting host phenotype will remain to be investigated [11].

### Biotic Interactions and EPN Evolution

The evolution potential of EPNs and their associated microbiota might not solely be impacted by environmental stressors like temperature but might also be profoundly influenced by biotic interactions within their microbial communities. Experimental passaging of microbial populations—whether in isolation or co-culture—has revealed how interspecies dynamics can either constrain or promote evolutionary adaptation [48]. For example, studies on soil bacterial communities demonstrate that the presence of competitors can limit abiotic adaptation. When evolved alone, *Pseudomonas fluorescens* exhibited higher fitness than when co-evolved with *Pseudomonas putida*, due to mutations that conferred a growth advantage in soil—an advantage that vanished in the presence of *P. putida* [48].

In the context of EPNs, these findings suggest that the evolutionary trajectory of their microbiota can be shaped not only by temperature but also by the competitive or cooperative dynamics among bacterial species. In our study, no significant change in microbiota composition could be observed between the two temperature treatments after only one cycle. However, the striking finding that significant shifts in microbiota composition were observed after 20 cycles at high temperatures, aligns with the above mentioned perspective, potentially reflecting indirect effects mediated by interspecies interactions depending on thermal regime. The remarkable enrichment of *Alcaligenes faecalis* in our experiment at 28 °C, barely detectable in samples at 23°C (Supp data S3), coincided with a reduction in the reproductive success of the entomopathogenic nematodes. Notably, *A. faecalis* has been previously reported to exert deleterious effects on the development and morphology of *Heterorhabditis* nematodes in culture conditions [49], including morphological abnormalities and *A. faecalis* observed under our experimental evolution conditions may impact nematode reproductive success. Interestingly, the enrichment of *Alcaligenes faecalis* was concomitent with a decrease in abundance of *Pseudomonas* and *Xenorhabdus* (Fig 5B), two bacterial species with entomopathogenic properties. However, in our conditions, temperature was not responsible in modifying the EPNs’ insects killing ability, which remained high whatever the experimental conditions used. Our study in *S. carpocapsae* suggests that virulence and reproductive success are decoupled traits, one being unaffected and the other impaired by evolution at high temperature, contrary to what has been suggested in *Steinernema anomaly* [50]. The direct contribution of the bacterial microbiota to these phenotypic traits will remain to be deciphered. Our findings, combined with insights from other systems, indicate that temperature-driven shifts in microbiota composition is concomitant with impaired EPN performance, suggesting an interplay between abiotic stress and biotic interactions. Reshaping microbial communities (an approach also called microbiome engineering) might be a way to confirm a putative microbiota composition and EPN fitness correlation. Such approach could later open the way to bolster EPN performance during abiotic stress [51]. Studies on soil microbiomes indicate highly variable effects of global change factors (warming, elevated CO2, drought etc) on microbial alpha diversity, but suggest that rare microorganisms are more strongly affected by such factors than the dominant taxa [52, 53]. Future work should explore whether specific bacterial taxa within the EPN microbiota, including the rare ones, act as key species, facilitating or hindering adaptation to thermal stress.

Over long timescales, mutations may emerge and get fixed within the microbiota population (including in the symbiont *Xenorhabdus*, as observed after a few hundred generations [14, 38]), further influencing the adaptive landscape of the EPN’s holobiont. Thus, microbiome species may evolve and adapt to elevated temperatures and the new interaction networks, triggering further compositional changes through eco-evolutionary feedback. This process can ultimately help the nematode-bacteria complex adapt to high-temperature conditions.

### EPNs as Biocontrol agents in a Changing Climate

The broader implications of this work suggest that as global temperatures rise, the efficacy of EPN-based biocontrol may depend on our ability to predict and manipulate their evolutionary responses. In a context of climate change, future research could expand experimental evolution studies to include different and combined abiotic stressors [54], but also a broader range of EPN species and their microbiota, or various insect hosts, aiming at identifying universal and species-specific adaptive strategies, including among their microbiota. Ultimately, further studies may explore applications of EPN strains in agroecosystems with fluctuating thermal environments, assessing how microbiota-mediated adaptation translates to biocontrol efficacy.

## Conclusion

This study represents a critical first step in determining whether temperature is associated with differences in EPN bacterial microbiota composition during reproduction in insect hosts, providing insights into how this abiotic factor shapes the evolution of EPN life-history traits in controlled environments. While our experimental evolution approach did not allow us to study adaptation to elevated temperature, it revealed that both microbiota composition and reproductive success are altered through successive cycles—especially under sub-optimal high temperatures—and triggered questions on the population genetics parameters of these populations. The microbiota composition was modified between the two experimental temperatures, concomitantly to the fitness decline, yet the contributions of the broader microbiota in determining EPN life-history traits remain largely underexplored [3]. These findings open new paths for exploring whether microbial changes directly influence EPN fitness or are simply a consequence of thermal regimes, with implications for predicting EPN adaptation to fluctuating environmental conditions in a climate change context, which is crucial for sustainable insect pest management.

## Acknowledgements

This research was funded by INRAE-SPE and UM-CLAPAS. We also thank Raphaël Bousquet for technical assistance with nematodes characterization.

## Author contributions

JB and SB designed the experiments; CR, NF, JCO and JB performed the experiments; CR, NF, JCO, JB and SB analyzed the data. JB drafted the manuscript. JB, SB and JCO reviewed and edited the manuscript. All authors have approved its final version.

## Additional Information

The authors declare no conflict of interest.

