## Supplemental Data (Figures) for "Experimental evolution at high temperature in a Lepidopteran host of an entomopathogenic nematode and its associated microbiota"

**Supp Data S1**. Design of the serial passages experiment

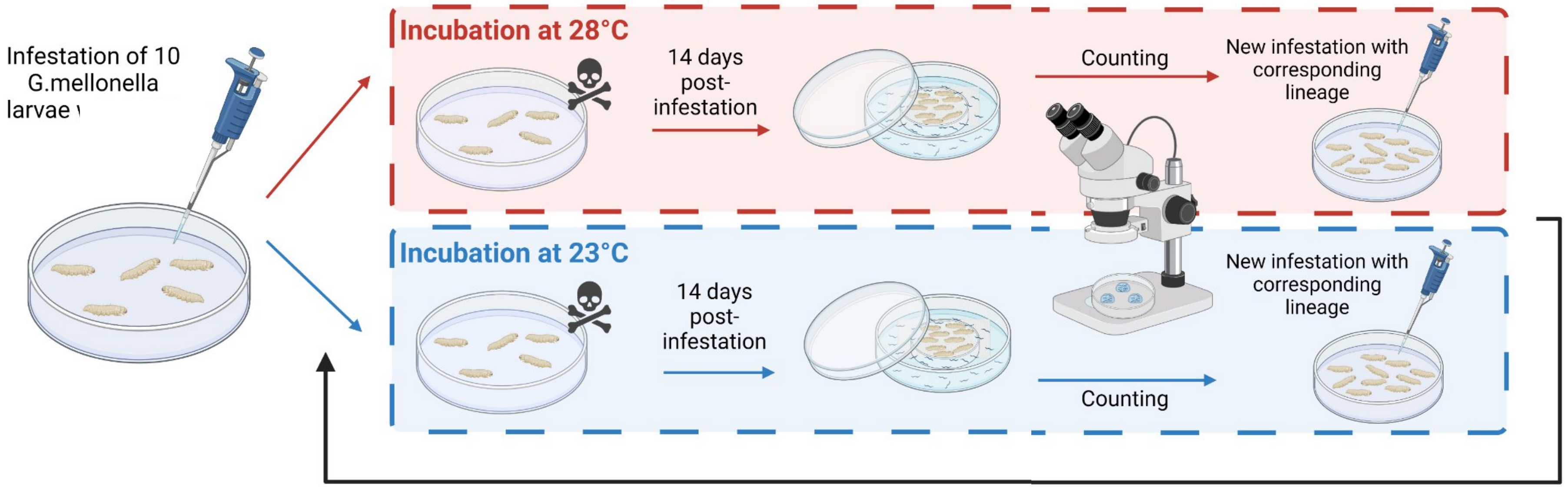


12 individually evolved lineages
- 6 incubated at 23°C = EE23
- 6 incubated at 28°C = EE28

#### Supp Data S2

Cumulated data of the Reproductive success of the evolved lineages

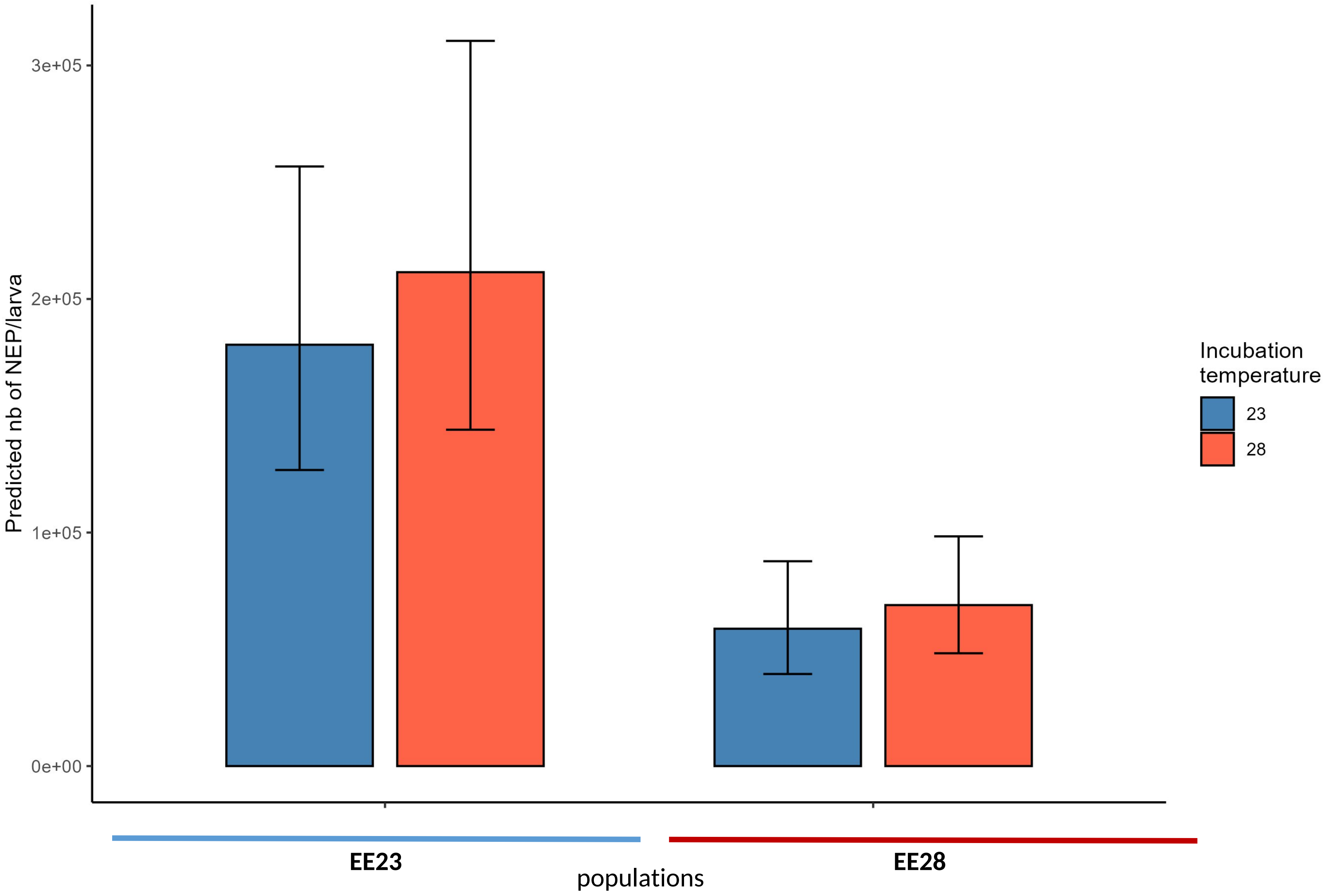


*Galleria mellonella* larvae were infested by 100 *S. carpocapsae* IJs, from 6 evolved lineages after 20 serial passages at 23°C (EE23) or 6 evolved lineages at 28 °C (EE28), and incubated at either 23°C (blue) or 28°C (red). The dead larvae were put on a White-trap and incubated at either 23°C (blue) or 28°C (red). IJs emerging from *G. mellonella* cadavers were harvested at various days post-infestation (dpi) as presented in Fig. 3. The number of cumulated IJs was monitored (i.e., all the IJs that emerged after 28 dpi). Each experiment contained 10 larvae per lineage/temperature (totalizing 60 larvae for each of the presented histogram).

### SuppData S4

Rarefaction curves

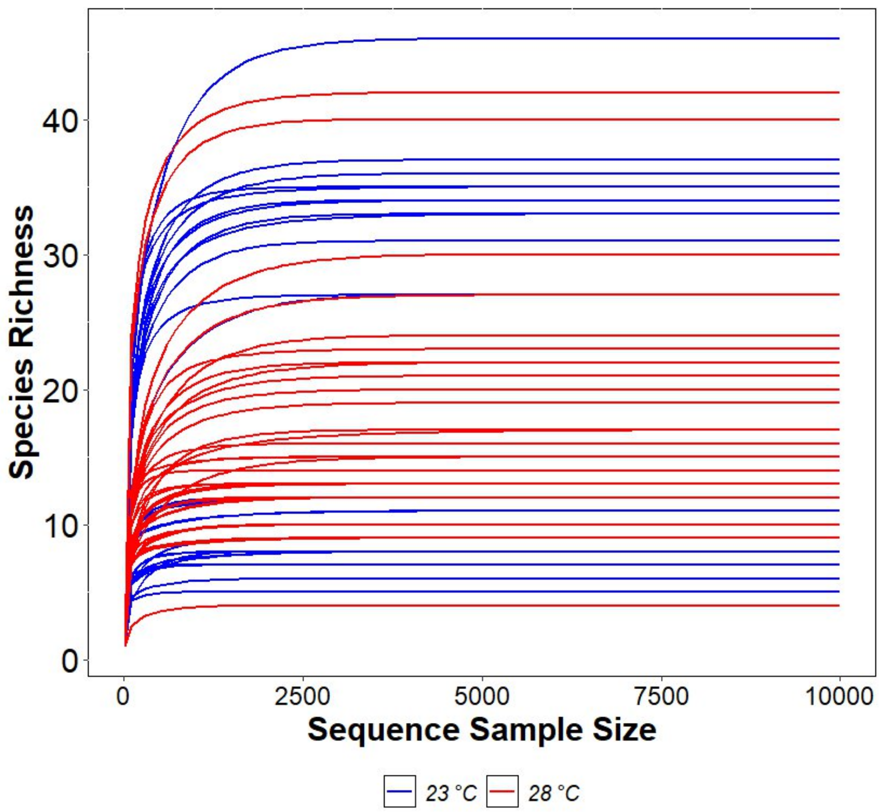


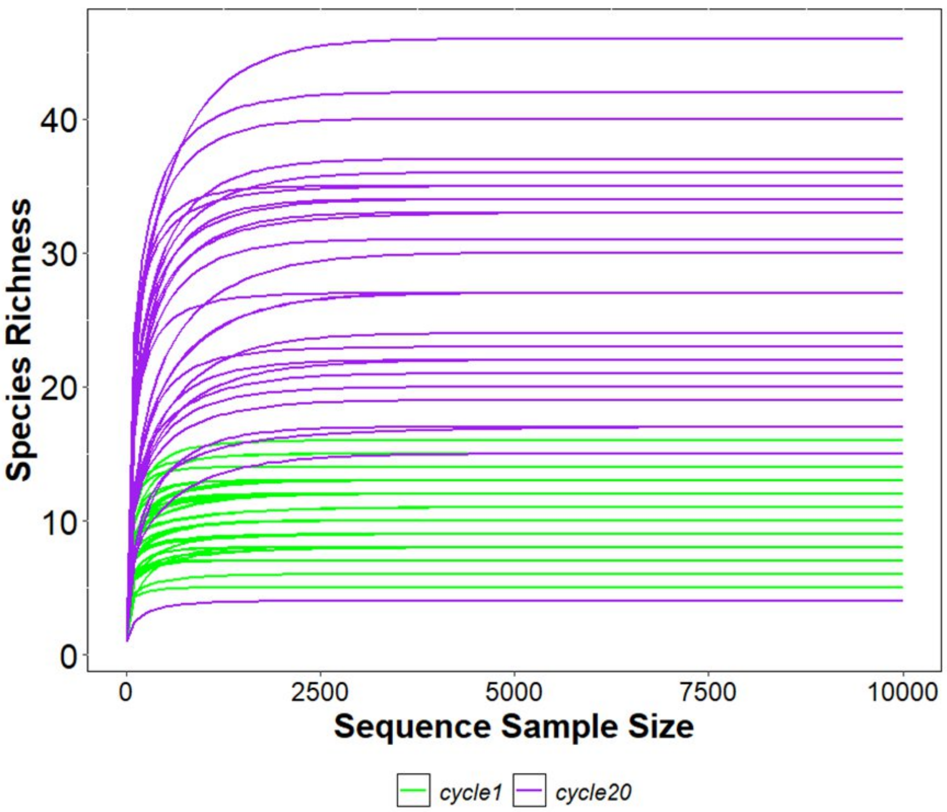


**Figure Legend: Rarefaction curves.** For each sample, species richness was plotted as a function of sequencing depth, with the number of observed ASVs on the y-axis and the number of sequences on the x-axis. Samples were colored according to incubation temperature (23°C and 28°C) or to cycle (cycle 1 and cycle 20).

### Supp data Fig. S5

## A.

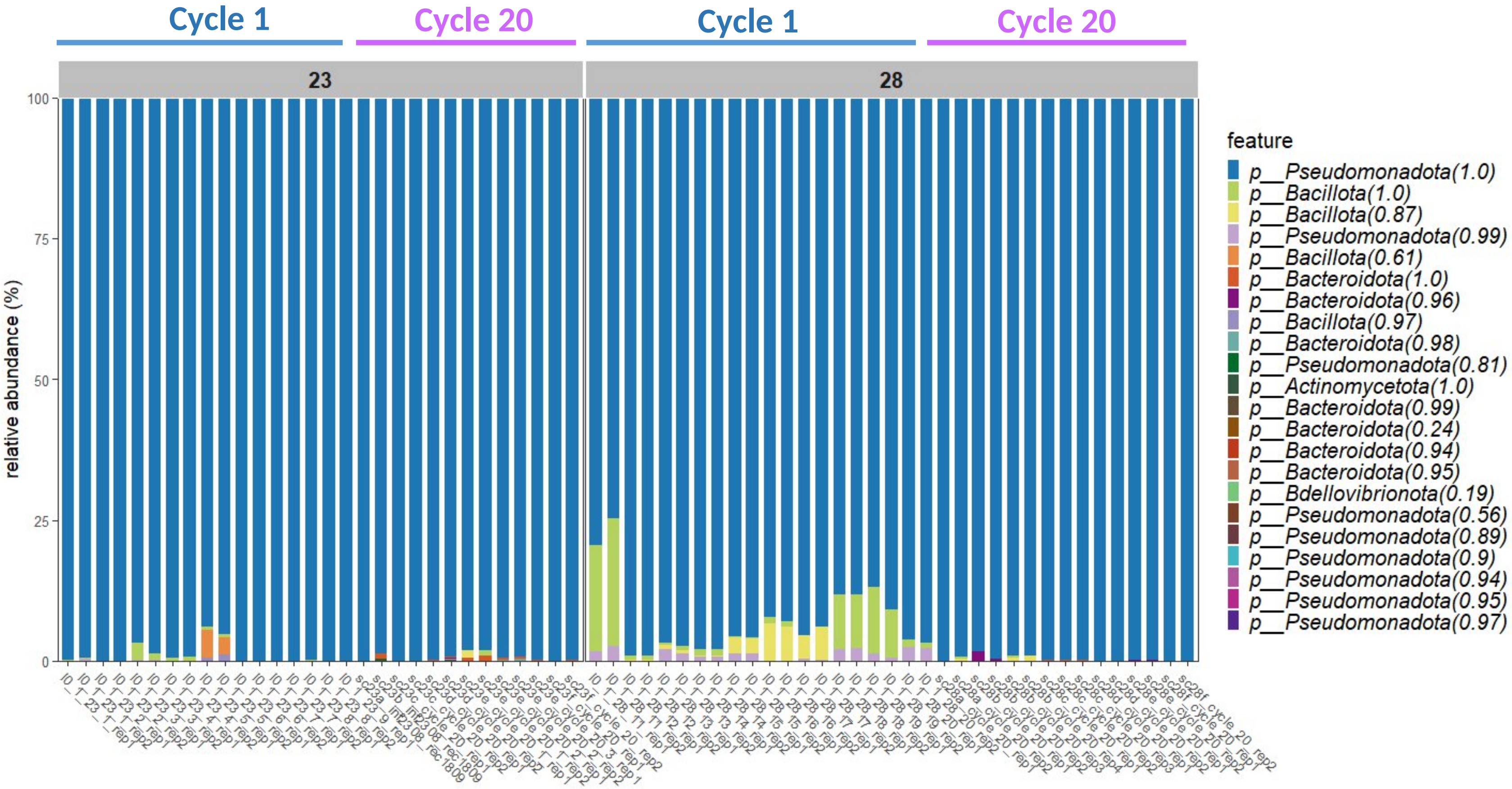

### Supp data Fig. S5

## B.

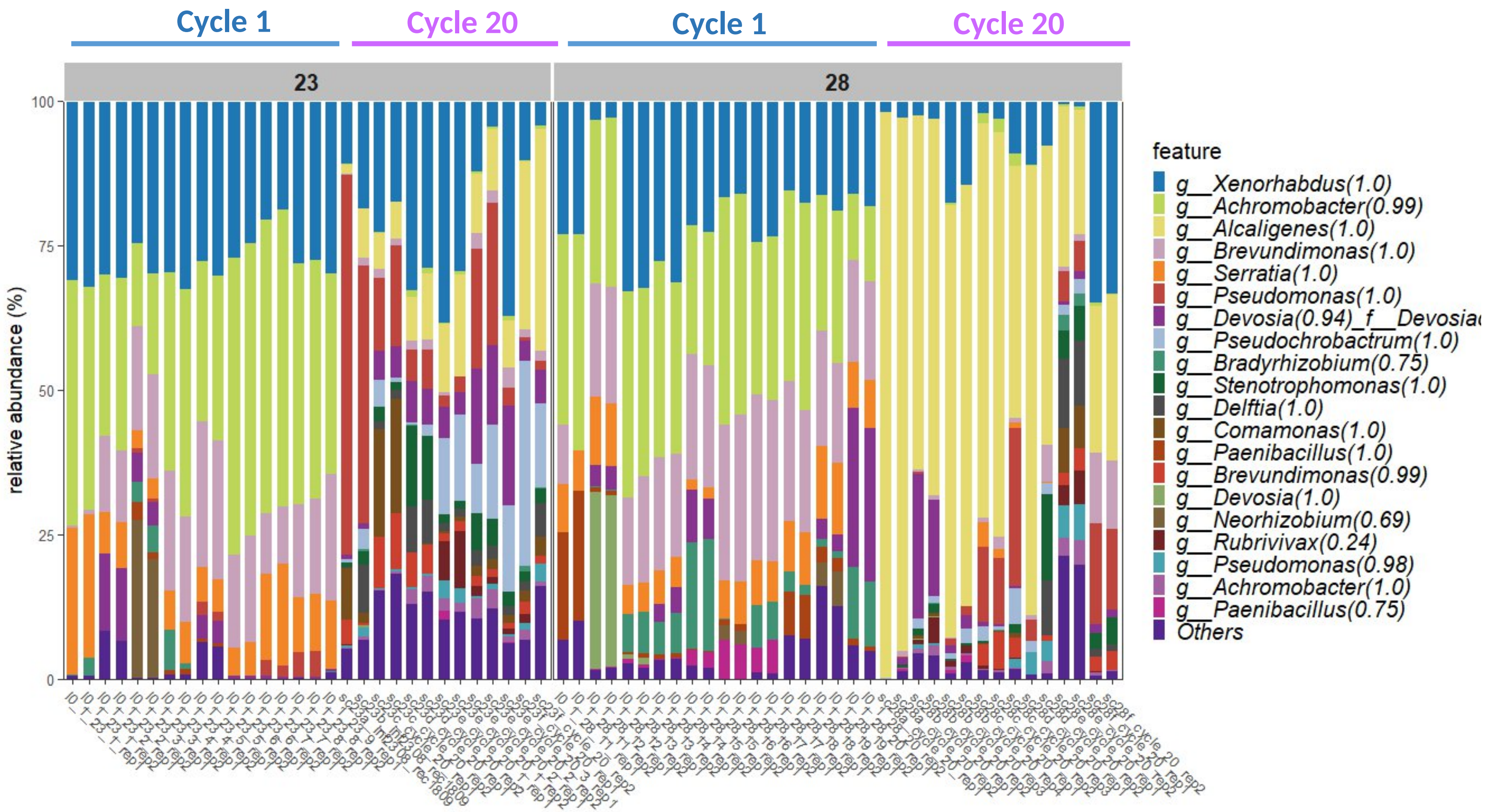

### Supp data Fig. S5

## C.

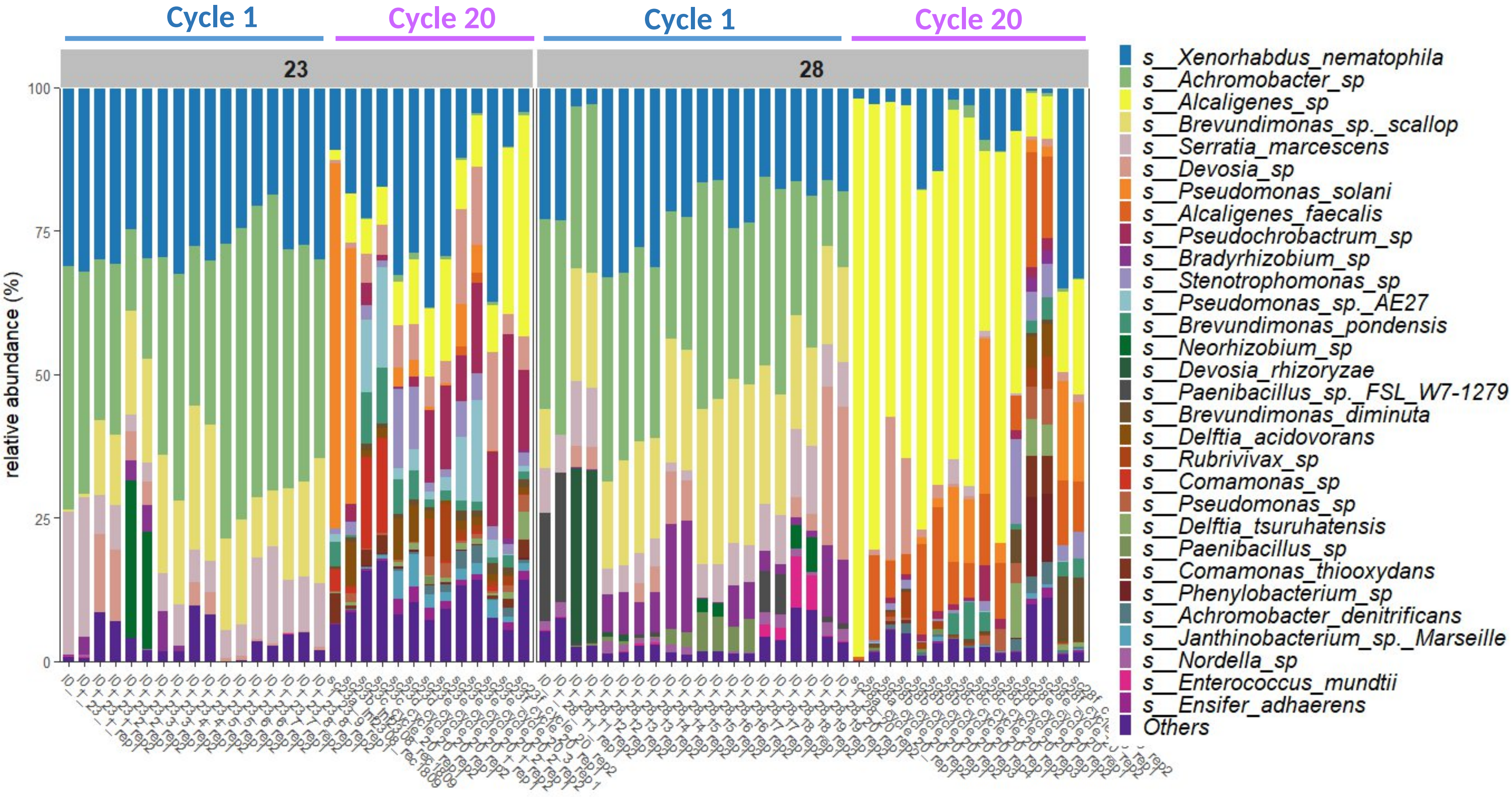
