## Supplemental Data (Table) for "Experimental evolution at high temperature in a Lepidopteran host of an entomopathogenic nematode and its associated microbiota"

| ASVs | Taxonomy (RDD classifier) | observation_ | sc28a_cycle_2 | sc28a_cycle_ | sc28b_cycle_ | sc28b_cycle_ | sc28b_cycle_ | sc28b_cycle_ | sc28c_cycle_ | sc28c_cycle_2 | sc28c_cycle_2 | sc23e_cycle_ | sc23e_cycle_ | sc23e_cycle_ | sc23e_cycle_ | sc23e_cycle_ | sc23b_inf230 |
|---|---|---|---|---|---|---|---|---|---|---|---|---|---|---|---|---|---|
| ASV_1 | Bacteria(1.0);Pseudomonadota(1.0);Gammaproteobacteria(1.0);Enterobacte | 412911 | 850 | 781 | 365 | 1343 | 4642 | 2701 | 980 | 1414 | 2519 | 6817 | 1101 | 4491 | 1471 | 14298 | 8477 |
| ASV_3 | Bacteria(1.0);Pseudomonadota(1.0);Betaproteobacteria(1.0);Burkholderiales | 377924 | 0 | 0 | 0 | 0 | 0 | 0 | 840 | 986 | 578 | 5 | 1 | 0 | 0 | 0 | 40 |
| ASV_6 | Bacteria(1.0);Pseudomonadota(1.0);Alphaproteobacteria(1.0);Caulobacterale | 203919 | 0 | 0 | 0 | 0 | 0 | 0 | 0 | 0 | 0 | 0 | 0 | 0 | 0 | 0 | 0 |
| ASV_2 | Bacteria(1.0);Pseudomonadota(1.0);Betaproteobacteria(1.0);Burkholderiales | 169023 | 125 | 117 | 7355 | 26775 | 15158 | 9911 | 28695 | 28565 | 8371 | 16 | 1 | 29 | 19 | 19 | 32 |
| ASV_9 | Bacteria(1.0);Pseudomonadota(1.0);Gammaproteobacteria(1.0);Enterobacte | 109398 | 0 | 0 | 0 | 0 | 0 | 0 | 2045 | 807 | 254 | 0 | 0 | 0 | 0 | 0 | 0 |
| ASV_5 | Bacteria(1.0);Pseudomonadota(1.0);Gammaproteobacteria(1.0);Pseudomona | 81912 | 0 | 0 | 13 | 0 | 281 | 286 | 6092 | 4887 | 7438 | 97 | 16 | 2463 | 1447 | 42 | 20532 |
| ASV_4 | Bacteria(1.0);Pseudomonadota(1.0);Betaproteobacteria(1.0);Burkholderiales | 63274 | 41123 | 19256 | 0 | 0 | 0 | 0 | 5 | 14 | 3 | 18 | 5 | 116 | 49 | 25 | 50 |
| ASV_7 | Bacteria(1.0);Pseudomonadota(1.0);Betaproteobacteria(1.0);Burkholderiales | 48762 | 0 | 0 | 1 | 13 | 6 | 2 | 43 | 78 | 13 | 2048 | 654 | 2903 | 2717 | 3064 | 3532 |
| ASV_12 | Bacteria(1.0);Pseudomonadota(1.0);Alphaproteobacteria(1.0);Hyphomicrobi | 41502 | 57 | 0 | 146 | 300 | 266 | 247 | 479 | 203 | 1717 | 2242 | 546 | 2892 | 5049 | 4890 | 1081 |
| ASV_18 | Bacteria(1.0);Pseudomonadota(1.0);Betaproteobacteria(1.0);Burkholderiales | 39482 | 151 | 3214 | 797 | 1286 | 3791 | 3121 | 2825 | 1967 | 3205 | 0 | 0 | 535 | 540 | 25 | 0 |
| ASV_31 | Bacteria(1.0);Pseudomonadota(1.0);Alphaproteobacteria(1.0);Hyphomicrobi | 37281 | 0 | 0 | 0 | 0 | 0 | 0 | 0 | 0 | 0 | 0 | 0 | 0 | 0 | 0 | 1 |
| ASV_38 | Bacteria(1.0);Pseudomonadota(0.98);Alphaproteobacteria(0.98);Hyphomicro | 31040 | 0 | 0 | 0 | 0 | 0 | 0 | 0 | 0 | 0 | 0 | 0 | 0 | 0 | 0 | 0 |
| ASV_23 | Bacteria(1.0);Pseudomonadota(1.0);Alphaproteobacteria(1.0);Hyphomicrobi | 29224 | 8 | 279 | 3403 | 7273 | 369 | 427 | 337 | 234 | 125 | 688 | 82 | 1527 | 907 | 4246 | 275 |
| ASV_20 | Bacteria(1.0);Pseudomonadota(1.0);Alphaproteobacteria(1.0);Caulobacterale | 24336 | 0 | 0 | 5 | 10 | 0 | 0 | 1814 | 2982 | 951 | 248 | 68 | 662 | 443 | 393 | 208 |
| ASV_51 | Bacteria(1.0);Pseudomonadota(1.0);Alphaproteobacteria(1.0);Hyphomicrobi | 21983 | 0 | 0 | 0 | 0 | 0 | 0 | 0 | 0 | 0 | 0 | 0 | 0 | 0 | 0 | 0 |
| ASV_57 | Bacteria(1.0);Bacillota(1.0);Bacilli(1.0);Bacillales(1.0);Paenibacillaceae(1.0);Paer | 19888 | 0 | 0 | 0 | 0 | 0 | 0 | 0 | 0 | 0 | 0 | 0 | 0 | 0 | 0 | 0 |
| ASV_34 | Bacteria(1.0);Pseudomonadota(1.0);Alphaproteobacteria(1.0);Caulobacterale | 17016 | 11 | 177 | 9 | 55 | 17 | 15 | 79 | 355 | 129 | 89 | 0 | 596 | 372 | 696 | 398 |
| ASV_40 | Bacteria(1.0);Pseudomonadota(1.0);Gammaproteobacteria(1.0);Xanthomon | 15583 | 6 | 27 | 5 | 58 | 30 | 51 | 81 | 2 | 16 | 200 | 50 | 2341 | 1436 | 881 | 0 |
| ASV_64 | Bacteria(1.0);Pseudomonadota(1.0);Betaproteobacteria(1.0);Burkholderiales | 15342 | 0 | 0 | 0 | 0 | 0 | 0 | 2 | 6 | 2 | 0 | 0 | 0 | 0 | 0 | 1 |
| ASV_61 | Bacteria(1.0);Pseudomonadota(1.0);Alphaproteobacteria(1.0);Hyphomicrobi | 14469 | 0 | 0 | 0 | 0 | 0 | 0 | 0 | 0 | 0 | 0 | 0 | 0 | 0 | 0 | 0 |
| ASV_21 | Bacteria(1.0);Pseudomonadota(1.0);Betaproteobacteria(1.0);Burkholderiales | 12756 | 0 | 32 | 6 | 3 | 27 | 6 | 34 | 25 | 15 | 229 | 23 | 563 | 452 | 458 | 3254 |
| ASV_28 | Bacteria(1.0);Pseudomonadota(1.0);Gammaproteobacteria(1.0);Xanthomon | 12308 | 0 | 84 | 153 | 605 | 41 | 29 | 98 | 24 | 323 | 63 | 2 | 23 | 46 | 49 | 920 |
| ASV_30 | Bacteria(1.0);Pseudomonadota(1.0);Gammaproteobacteria(1.0);Pseudomon | 11722 | 0 | 33 | 0 | 0 | 11 | 0 | 5 | 20 | 37 | 178 | 70 | 3875 | 5544 | 610 | 0 |
| ASV_41 | Bacteria(1.0);Pseudomonadota(1.0);Betaproteobacteria(1.0);Burkholderiales | 11621 | 0 | 0 | 0 | 0 | 0 | 0 | 0 | 0 | 0 | 1 | 14 | 53 | 82 | 267 | 364 |
| ASV_17 | Bacteria(1.0);Pseudomonadota(1.0);Gammaproteobacteria(1.0);Pseudomon | 10712 | 0 | 0 | 0 | 0 | 0 | 0 | 0 | 0 | 0 | 80 | 17 | 42 | 59 | 525 | 1 |
| ASV_54 | Bacteria(1.0);Pseudomonadota(1.0);Betaproteobacteria(1.0);Burkholderiales | 10233 | 0 | 0 | 8 | 15 | 0 | 0 | 15 | 12 | 27 | 32 | 30 | 381 | 307 | 149 | 244 |
| ASV_16 | Bacteria(1.0);Pseudomonadota(1.0);Betaproteobacteria(1.0);Burkholderiales | 8774 | 0 | 0 | 118 | 1921 | 272 | 252 | 228 | 99 | 75 | 1190 | 370 | 640 | 398 | 353 | 6 |
| ASV_42 | Bacteria(1.0);Pseudomonadota(1.0);Betaproteobacteria(1.0);Burkholderiales | 7960 | 0 | 0 | 0 | 0 | 11 | 151 | 0 | 5 | 53 | 425 | 53 | 697 | 612 | 1167 | 14 |
| ASV_85 | Bacteria(1.0);Pseudomonadota(0.99);Alphaproteobacteria(0.99);Hyphomicro | 7458 | 0 | 0 | 0 | 0 | 0 | 0 | 0 | 0 | 0 | 0 | 0 | 0 | 0 | 0 | 0 |
| ASV_94 | Bacteria(1.0);Pseudomonadota(1.0);Alphaproteobacteria(1.0);Caulobacterale | 7348 | 0 | 0 | 0 | 0 | 0 | 0 | 0 | 0 | 0 | 0 | 0 | 0 | 0 | 0 | 0 |
| ASV_96 | Bacteria(1.0);Bacillota(1.0);Bacilli(1.0);Lactobacillales(1.0);Enterococcaceae(1.0 | 7322 | 0 | 0 | 0 | 16 | 0 | 0 | 0 | 0 | 0 | 0 | 0 | 0 | 0 | 14 | 0 |
| ASV_37 | Bacteria(1.0);Pseudomonadota(1.0);Betaproteobacteria(1.0);Burkholderiales | 7207 | 0 | 38 | 98 | 675 | 124 | 70 | 85 | 158 | 52 | 268 | 54 | 1153 | 953 | 83 | 67 |
| ASV_49 | Bacteria(1.0);Pseudomonadota(1.0);Alphaproteobacteria(1.0);Caulobacterale | 6891 | 0 | 1 | 10 | 20 | 0 | 13 | 43 | 15 | 4 | 9 | 5 | 71 | 27 | 28 | 31 |
| ASV_29 | Bacteria(1.0);Pseudomonadota(1.0);Alphaproteobacteria(1.0);Hyphomicrobi | 6795 | 0 | 0 | 0 | 0 | 0 | 0 | 0 | 0 | 0 | 54 | 0 | 2807 | 2124 | 170 | 137 |
| ASV_99 | Bacteria(1.0);Pseudomonadota(1.0);Gammaproteobacteria(1.0);Pseudomon | 6621 | 0 | 0 | 0 | 0 | 0 | 0 | 0 | 0 | 0 | 0 | 0 | 0 | 0 | 0 | 0 |
| ASV_74 | Bacteria(1.0);Pseudomonadota(1.0);Alphaproteobacteria(1.0);Hyphomicrobi | 5619 | 0 | 49 | 18 | 5 | 0 | 41 | 4 | 4 | 12 | 385 | 51 | 342 | 345 | 60 | 151 |
| ASV_105 | Bacteria(1.0);Bacillota(0.87);Bacilli(0.87);Bacillales(0.86);Paenibacillaceae(0.78); | 5583 | 0 | 0 | 12 | 0 | 3 | 5 | 0 | 0 | 0 | 0 | 0 | 0 | 0 | 0 | 0 |
| [illegible] | | | | | | | | | | | | | | | | | |
